# Quantitative Theory for the Diffusive Dynamics of Liquid Condensates

**DOI:** 10.1101/2021.03.08.434288

**Authors:** Lars Hubatsch, Louise M. Jawerth, Celina Love, Jonathan Bauermann, T.-Y. Dora Tang, Stefano Bo, Anthony A. Hyman, Christoph A. Weber

## Abstract

To unravel the biological functions of membraneless liquid condensates it is crucial to develop a quantitative understanding of the physics underlying their dynamics. Key processes within such condensates are diffusion and material exchange with their environment. Experimentally, diffusive dynamics are typically probed via fluorescent labels. However, to date we lack a physics-based quantitative framework for the dynamics of labeled condensate components. Here, we derive the corresponding theory, building on the physics of phase separation, and quantitatively validate this framework via experiments. We show that using our theory we can precisely determine diffusion coefficients inside liquid condensates via a spatio-temporal analysis of fluorescence recovery after photobleaching (FRAP) experiments. We showcase the accuracy and precision of our approach by considering space- and time-resolved data of protein condensates and two different polyelectrolyte-coacervate systems. Strikingly, our theory can also be used to determine the diffusion coefficient in the dilute phase and the partition coefficient, without relying on fluorescence measurements in the dilute phase. This bypasses recently described quenching artefacts in the dense phase, which can underestimate partition coefficients by orders of magnitude. Our experimentally verified theory opens new avenues for theoretically describing molecule dynamics in condensates, measuring concentrations based on the dynamics of fluorescence intensities and quantifying rates of biochemical reactions in liquid condensates.

## Introduction

Liquid phase separation has emerged as an organizing principle in biology and is thought to underlie the formation of various membrane-less cellular organelles (***Banani et al. (2017)***). Hallmark properties of such organelles are their rapid formation and dissolution, their fusion, and their wetting to membranes (***Hyman et al. (2014)***). Moreover, phase-separated organelles exchange material with their environment leading to dynamic sequestration of molecules, which affects biochemical processes by spatial redistribution of reactants (***Moon et al. (2019)***; ***Lyon et al. (2020)***; ***Saha et al. (2016)***; ***Guillén-Boixet et al. (2020)***; ***Sanders et al. (2020)***; ***Yang et al. (2020)***). Probing the dynamics of condensate components is thus imperative for a quantitative understanding of how they affect the cellular biochemistry (***Mir et al. (2019)***).

To probe the dynamics of condensates, biomolecules are typically labelled with fluorescent tags. In general, in systems with tagged molecules, various methods exist to characterize molecular properties such as binding rates and diffusion coefficients, including fluorescence correlation spectroscopy (FCS) (***Ries and Schwille (2012)***; ***Rigler and Elson (2012)***, single-particle tracking (SPT) (***Tinevez et al. (2017)***; ***Saxton and Jacobson (1997)***), and fluorescence recovery after photobleaching (FRAP) (***Diaspro (2010)***; ***Stasevich et al. (2010)***). However, interpretation of the experimental data acquired from such methods requires a rigorous derivation accounting for the underlying physicochemcial processes. This derivation has been achieved for some biological systems and processes, but is lacking for condensates formed by liquid phase separation. Processes that are well-understood include membrane-cytoplasmic exchange and transport (***Sprague et al. (2004)***; ***Robin et al. (2014)***; ***Goehring et al. (2010)***) as well as chemical reactions (***Elson (2001)***) or filament turnover (***McCall et al. (2019)***). For liquid condensates, various phenomenological fit functions have been proposed in the literature (e.g. ***Patel et al. (2015)***; ***Banerjee et al. (2017***); ***Hubstenberger et al. (2013)***, for a broader summary see ***Taylor et al. (2019)***). However, it was recently shown that these fits lead to wildly differing estimates of the diffusion constant inside, ***D**_in_* (***Taylor et al. (2019)***). Taylor et al. showed that these discrepancies were attributed to unrealistic assumptions, e.g. infinitely large droplets or infinitely fast diffusion outside the bleach area. However, a conclusive answer of what governs the dynamics of molecules across a condensate interface could not be reached.

Here, we first introduce a quantitative FRAP method to extract the diffusion coefficient inside ***D***_in_, purely based on fluorescence measurements inside droplets, without resorting to unrealistic assumptions or requiring knowledge about the partition coefficient, ***P*** or diffusion outside, ***D***_out_. Using non-equilibrium thermodynamics, we then derive the theory that connects dynamics inside and outside of the droplet via transport across a finite interface. We show that this theory fits our experimentally observed dynamics with striking agreement. Surprisingly, we theoretically show that it can be used to extract all relevant parameters of the system, ***P***, ***D***_in_ and ***D***_out_, purely based on knowledge of the dynamics inside the droplet. We show that our measurements are agnostic to breaking radial symmetry, for example by introducing a coverslip or neighbouring droplets. Our approach does not suffer from typical limitations of fluorescence-based concentration measurements, such as low fluorescence in the dilute phase and fluorophore quenching in the rich phase. We anticipate that this new understanding will open the door to characterizing dynamical properties such as chemical rates and rheological parameters in multi-component, phase-separated systems.

## Results

### Determining the diffusion constant inside liquid condensates

First, we discuss a quantitative method to extract diffusion coefficients of biomolecules in a condensate. After bleaching a liquid condensate, bleached molecules diffuse out and unbleached molecules diffuse into the condensate, until the unbleached components reach the spatially homogeneous level prior to bleaching (Fig. 1a). Inside a spherical condensate of radius ***R***, the concentration of unbleached components, *c_u_*(*r, t*), follows a diffusion equation (for derivation, see subsequent section),

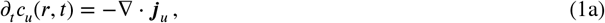

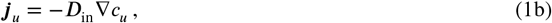

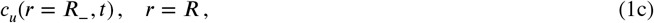

where *c_u_*(*r* = ***R***_, *t*) is the time-dependent concentration directly inside the interface at ***R*** = ***R***_. Here, ***R*** denotes the radial distance to the center of the condensate. The flux ***j**_u_* is given by Fick’s law (Eq. (1b)). It vanishes at the condensate center. Moreover, we have ∇ = ***e**_r_∂_r_*, with ***e_r_*** denoting the radial unit vector. During FRAP, the concentration at the interface, ***c_u_***(***r*** = ***R***_, *t*), changes with time (Fig.1b) and is determined by the physical properties of the condensate environment. This environment is characterized by the diffusion constant and the concentration of unbleached components outside, the distribution of neighbouring condensates as well as system boundaries like the coverslip.

**Figure 1.**
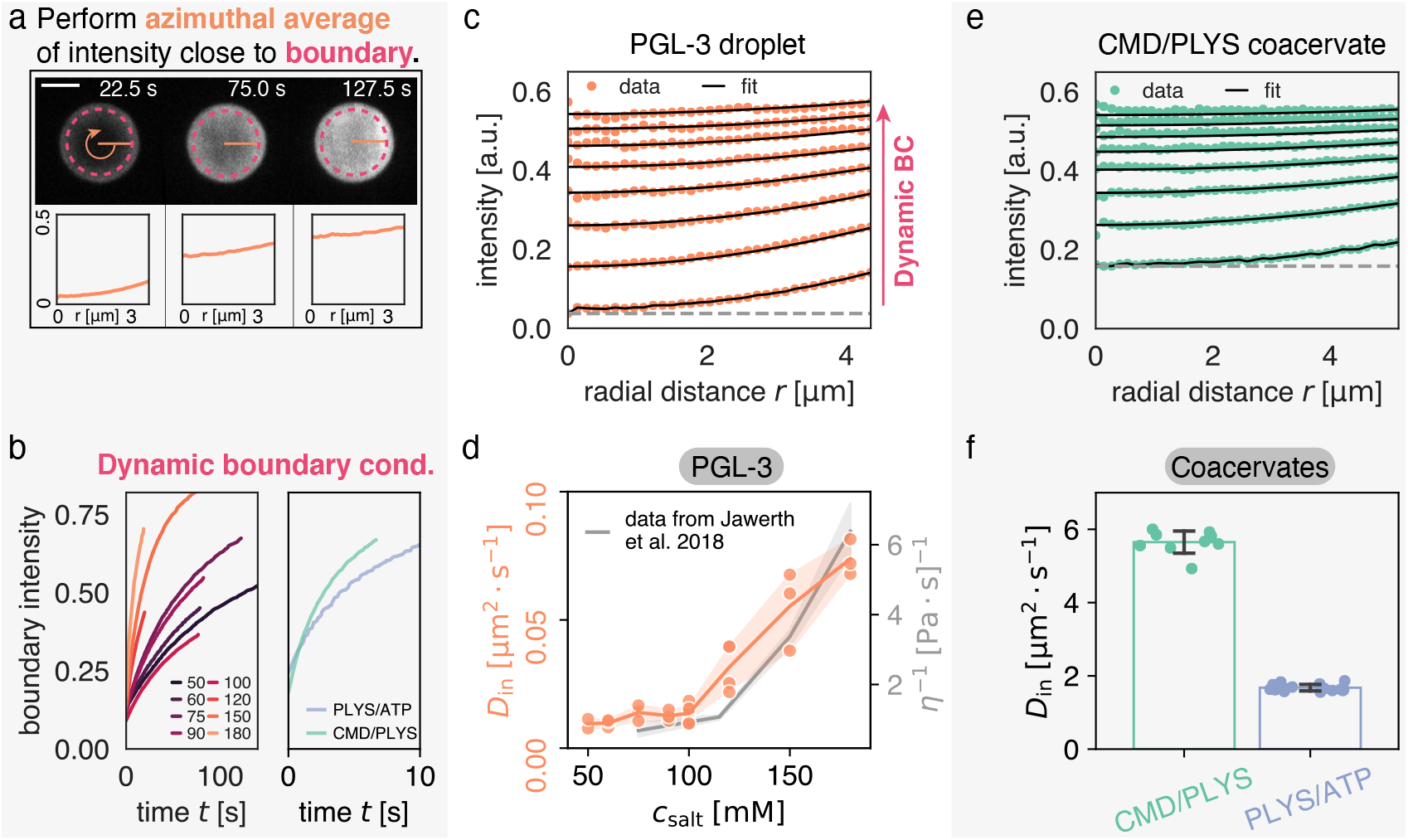
Quantitatively measuring ***D***_in_ by extracting concentration at condensate interface. (**a**) Time course of FRAP recovery after full bleach for PLYS/ATP coacervate droplet. Droplet interface (magenta) and azimuthal average (orange) are highlighted. Note, data close to the droplet boundary cannot be fit due to optical artefacts giving rise to an artificially broad interface. Thus, fluorescence intensity is extracted approximately 1.4 *μ*m away from the interface (see methods). Lower panels show azimuthal averages at different time points. Scale bar, 5 *μ*m. (**b**) Azimuthal average of fluorescence intensity inside condensates near the interface (magenta in (a)) for PGL-3 droplets at different salt concentrations (left, units: mM) and two coacervate systems (right). (**c**) Flurorescence recovery inside a PGL-3 condensate, along the radial direction (azimuthal averages, see (a)). ***D***_in_ is extracted by global fitting of Eq. 1b to the experimental profiles, using the experimentally extracted initial and boundary conditions (see panel (b)). The grey line indicates an offset that comes about due to incomplete bleaching, and a small but visible fast and uniform recovery with unknown cause (see Methods). (**d**) Comparison between ***D***_in_ and viscosity *η* for PGL-3 condensates at different salt concentrations. Viscosity data taken from (***Jawerth et al. (2018))*** for untagged PGL-3. Note, GFP-tagged PGL-3 has a higher viscosity than untagged PGL-3, which means the hydrodynamic radius of PGL-3:GFP cannot be directly computed from this panel (***Jawerth et al. (2020)).*** (**e**) Same as (c) but for a CMD/PLYS coacervate droplet. (**f**) Diffusion coefficients for coacervate systems. Each data point represents ***D***_in_ for a single droplet time course. Note the low spread of the measured values.

To initially bypass this complex dependence on the condensate environment, we propose to extract the fluorescence concentration of unbleached molecules directly inside of the condensate-bulk boundary *c_u_*(*r* = ***R***_,*t*) from experimental data after photobleaching (Fig.1a,b). Using this experimentally determined dynamic boundary condition, we can accurately determine the diffusion constant inside a condensate, ***D***_in_ (Fig.1d,f). Following this idea, we fit the solutions of Eq. (1) to spatio-temporal experimental data, with ***D***_in_ as the only fit parameter. We find excellent agreement between experimentally measured and fitted concentration profiles (Fig.1c,e and supp. movies 1,2,3). Specifically, we consider condensates composed of PGL-3, a main protein component of P granules in the *C. elegans* embryo (***Brangwynne et al. (2009); Griffin (2015))***, as well as two synthetic polyelectrolye-complex coacervate systems, Polylysine/ATP (PLYS/ATP) and Carboxymethyldextran/Polylysine (CMD/PLYS).

We first compared ***D***_in_ of PGL-3 for different salt concentrations between 50 mM and 180 mM (see Methods). We find that ***D***_in_ varies by roughly one order of magnitude, between 0.009 *μ*m^2^ s^-1^ and 0.070 *μ*m^2^ s^-1^. Our trend is in good agreement with reported measurements of the viscosity *η*, determined by active micro-rheology (***Jawerth et al. (2018))*** for untagged PGL-3 (Fig. 1 d). Using viscosity data for GFP-tagged PGL-3 (***Jawerth et al. (2020))***, we use the Stokes-Sutherland-Einstein relationship ***D***_in_ = *k*_B_***T***/(6*πaη*) to estimate the hydrodynamic radius of PGL-3:GFP, *a* = 1.5 nm (***Einstein (1905); Sutherland (1905); von Smoluchowski (1906))***. This estimate is consistent with the value reported in Ref. ***Liarzi and Epel (2005)***.

For the coacervate systems, we find for the diffusion coefficients inside ***D***_in_ = (1.68 ± 0.09) *μ*m^2^ s^-1^ for PLYS/ATP coacervates and ***D***_in_ = (5.65 ± 0.32) *μ*m^2^ s^-1^ for CMD/PLYS coacervates; see (Fig. 1f). The standard deviation of these measurements is low enough such that even a single measurement provides a good estimate of ***D***_in_. Interestingly, ***D***_in_ for the coacervate droplets is about 10 times smaller than the diffusion constant of the dilute polyelectrolytes ***D***_out_ (***Arrio-Dupont et al. (1996); Morga et al. (2019))***.

### Theory for the dynamics of labeled molecules in phase-separating systems

To understand the physical origin of the time-dependent concentration of unbleached components at the condensate interface and the phenomenological Eqs. (1) for the diffusion inside a condensate, we need a full theory for the diffusion inside, outside and across phase boundaries. Here, we derive such a theory for a system that can be described by a binary, incompressible mixture prior to photobleaching. This binary mixture is composed of condensate material and solvent. The condensate material has a concentration profile, 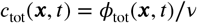, which can be expressed in terms of a volume fraction profile *ϕ*_tot_(***x***, *t*) by dividing through the molecular volume of the condensate material, *ν*. Due to incompressibility, the solvent volume fraction in such a binary mixture is given by (1 – *ϕ*_tot_). The system prior to photobleaching is assumed to be either at equilibrium, i.e., a single droplet and *∂_t_ϕ_tot_* = 0, or close to equilibrium, i.e., a system composed of many droplets undergoing slow Ostwald ripening and fusion and *∂_t_*ϕ*_tot_*) ≃ 0. Thus, the (quasi-) stationary profile *ϕ*_tot_(***x***) prescribes a physical constraint for FRAP dynamics.

After photobleaching, the system becomes a *ternary* incompressible mixture composed of bleached (*b*) and unbleached (*u*) components, as well as solvent (Fig. 2a). Introducing the volume fraction of the bleached and unbleached components, *ϕ_b_* and *ϕ_u_*, the physical constraint for the FRAP dynamics for a condensate at or close to equilibrium reads

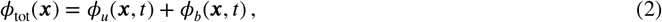

where the profiles of bleached and unbleached components depend on space and time during the FRAP dynamics, while *ϕ*_tot_(***x***) depends on space only. Immediately after photobleaching, unbleached molecules diffuse into the condensate leading to FRAP dynamics of unbleached molecules inside (Fig. 2b,c). At long times, the concentration profile of unbleached molecules approaches the profile prior to photobleaching, *ϕ*_tot_(***x***). The dynamics of both concentration profiles, bleached and unbleached molecules, *c_i_* = *ϕ_i_*/*ν_i_* (*i* = *b,u*), with *ν_i_* denoting the molecular volumes, is described by the following conservation laws (*j* = *u, b*),

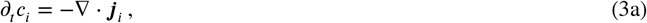

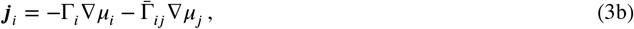

**Figure 2.**
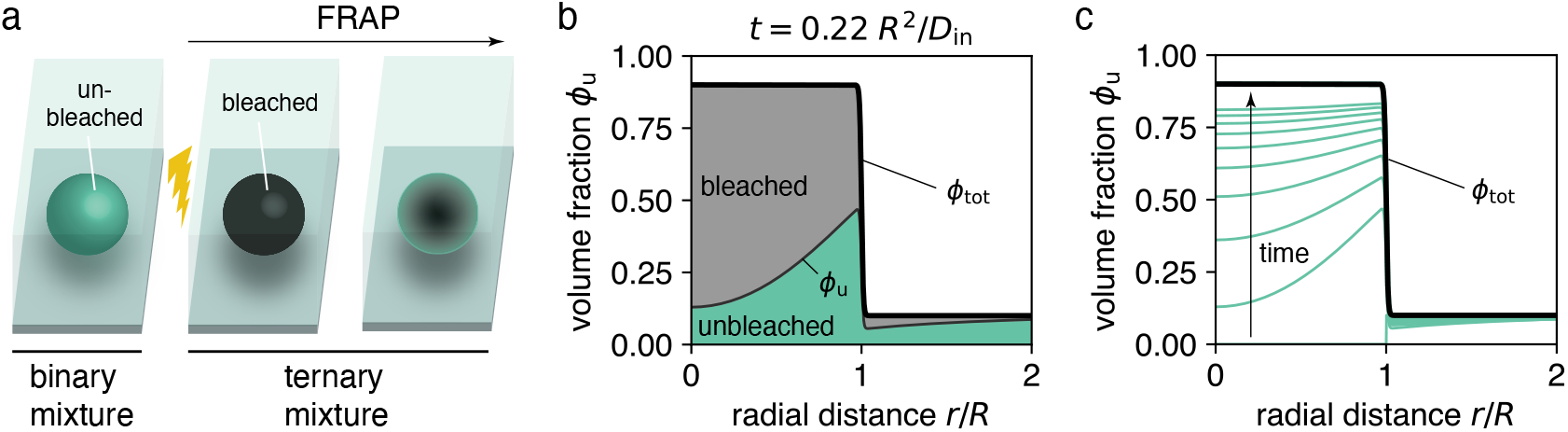
Ternary mixture accounts for the dynamics of bleached and unbleached molecules. (**a**) Before bleaching, a droplet that is composed of fluorescently labelled molecules can be described by a binary mixture, namely unbleached molecules and solvent. After bleaching, the system is composed of three components, bleached molecules, unbleached molecules and solvent. If the system was at equilibrium prior to bleaching, the sum of bleached and unbleached molecules forms a stationary, non-uniform profile *ϕ*_tot_(*r*) (see panel b). (**b**) Snapshot of model dynamics. Initial conditions are *ϕ*_u_(*r, t* = 0) = *ϕ*_out_ · Θ(*r* – ***R***), corresponding to a fully bleached droplet. Note that at any time we have *ϕ*_tot_ = *ϕ*_u_ + *ϕ*_b_· (**c**) Time course of spatial recovery. For long times, when nearly all bleached material has been exchanged, *ϕ*_u_ approaches *ϕ*_tot_. Panels (c,d) use radial symmetry for illustration purposes, however, the theory is general (see Fig. 3).

Here, Γ_*i*_ are the Onsager transport coefficients, often referred to as mobilities, and 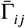 are the Onsager cross coupling coefficients obeying 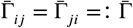. In general, both mobility coefficients depend on the volume fractions. To linear order, the flux ***J***_*i*_ is driven by gradients in exchange chemical potentials *μ_i_* and *μ_j_*. The exchange chemical potentials, *μ_i_* = *δ**F**/δ_c__i_*, are linked to the free energy, ***F*** = ∫ *d*^3^*x f*, where *f* denotes the free energy density. Expressing concentrations in terms of the volume fractions, *ϕ_i_* = *c_i_ν_i_*, we describe our incompressible ternary mixture after photobleaching by a Flory-Huggins free energy density (***Flory (1942); Huggins (1942); Krüger et al. (2018))***:

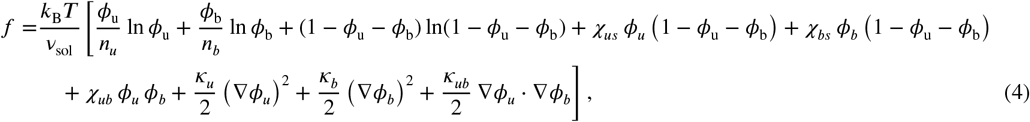

where we write the molecular volumes of bleached and unbleached components in *n_i_* multiples of the solvent molecular volume *ν*_sol_, i.e., *ν_i_* = *n_i_ν*_sol_. Moreover, *χ_ij_* denote dimensionless Flory-Huggins parameters characterizing the interactions between different components *i* and *j*, where subscript *s* indicates the solvent. The parameters *κ_i_* and *κ_ub_* characterize the free energy penalties for spatial heterogeneities and are linked to the surface tensions.

If photobleaching does not affect the molecular interactions or molecular volumes, the free energy density above can be simplified significantly (Appendix of Ref. (***Krüger et al. (2018))***. In this case, the interactions between unbleached and solvent, and bleached and solvent components are equal, *χ_us_* = *χ_bs_* =: *χ*, and cross interactions vanish to zero, *χ_ub_* = 0. Moreover, molecular volumes of bleached and unbleached components are equal, *n_u_* = *n_b_* =: *n* and the parameters characterizing free energy penalties for spatial heterogeneities obey *κ_u_* = *κ_b_* =: *κ* and *κ_ub_* = 2*κ*. Thus, the simplified free energy reads

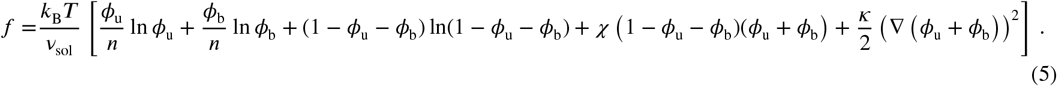

To ensure a constant diffusion coefficient in the dilute limits of the bleached and unbleached components, we employ the scaling ansatz for a ternary mixture, 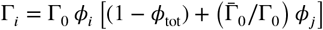 and 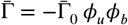. In general, both mobility functions, Γ_0_ and 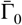, depend on the total volume fraction *ϕ*_tot_. For the limiting case where bleached and unbleached molecules are identical particles, we can choose 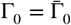. Applying the equilibrium FRAP condition (2) and using Eq. (3), we find that the concentration of unbleached components is governed by the following diffusion equation

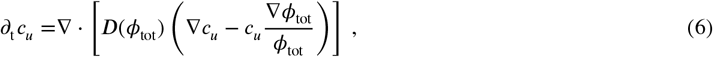

with a *ϕ*_tot_(*x*)-dependent diffusivity, ***D***(*ϕ*_tot_) = *k*_B_*T*Γ_0_(*ϕ*_tot_). As we show in Ref. (***Bo et al. (2021)***) a similar approach can be used to investigate single-molecule dynamics across phase boundaries.

Similar to Eq. (1), the diffusion equation above is linear in *c_u_*. However, the dynamics of unbleached components are affected by gradients in ***ϕ***_tot_(***x***) and components diffuse with different diffusion coefficients inside and outside the condensate, where in each phase ∇*ϕ*_tot_ = 0 (Fig. 2b). The position-dependence of *ϕ*_tot_(*x*) is given by the equilibrium condition of a homogeneous chemical potential of the binary mixture prior to photobleaching, which implies *∂*_t_*ϕ_tot_* = 0. For a radially symmetric system with ***R*** denoting the radial coordinate, 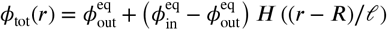, where 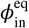 and 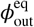 are the equilibrium volume fractions inside and outside, respectively, ***R*** is the droplet radius, and *ℓ* denotes the width of the interface. Moreover, *H* ((*r* – ***R***)/*ℓ*) is a function that decreases from one to zero at ***R*** = ***R*** on an interface width *ℓ*. For phase separation close to the critical point and large droplet sizes, *H*(*x*) = (1 + tanh (*x*))/2 (***Bray (1994); Weber et al. (2019))***. We numerically solve Eq. (6) using a finite element method (***Logg et al. (2012))*** in a finite domain of size ***L*** which is much larger than the droplet radius ***R*** and fit the solution to experimental data.

In summary, our model has seven parameters where four parameters, i.e., the equilibrium volume fractions 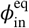 and 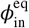, the interface width ŕ the droplet radius ***R***, characterize the equilibrium profile prior to bleaching *ϕ*_tot_(*x*). The remaining parameters are the system size ***L*** and the diffusion coefficients inside and outside, which are given as

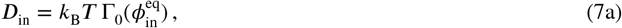

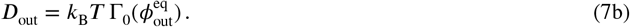

For the case of a single, spherical droplet with an infinitely thin interface (***Weber et al. (2019))***, we can derive an effective droplet model for the unbleached component from Eq. (6), where the dynamics of unbleached components inside and outside are given by diffusion equations that are coupled by boundary conditions (see App. 1 for the derivation):

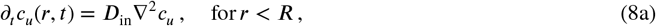

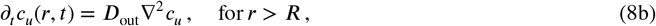

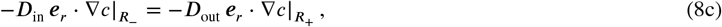

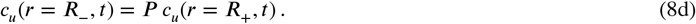

Here, ***R***_—_ and ***R***_+_ denote the radial position directly inside and outside the droplet interface, respectively. Eq. (8c) describes an equality of the fluxes directly inside and outside of the interface, respectively, and thereby expresses particle number conservation at the interface ***R*** = ***R***. Eq. (8d) describes a jump in concentration of unbleached components, which is determined by the thermodynamic partition coefficient

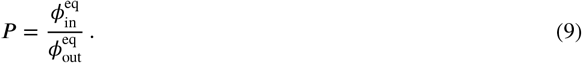

Moreover, the flux vanishes at the origin ***R*** = 0, *e_r_* · ∇*c*|_*r=0*_ = 0, and at the system boundary ***R*** = ***L***, *e_r_* · ∇ *c*|_*r=L*_ = 0. Eqs. (8) were phenomenologically proposed in Ref. (***Münchow et al. (2008))*** to study protein diffusion across the interface in aqueous two-Phase systems. Moreover, Eqs. (8) were also studied in the context of FRAP of protein condensates in Ref. (***Taylor et al. (2019))***, where ***D***_in_, ***D***_out_ and *P* were considered to be independent parameters. Strictly speaking, due to phase separation, the diffusion coefficients ***D***_in_ and ***D***_out_ are not independent which is evident in Eqs. (7). For example, in the absence of phase separation or at the critical point, 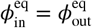 (i.e., *P* = 1), the diffusion coefficients inside and outside must be equal, ***D***_in_ = ***D***_out_. Specifically, Eqs. (7) can be written as

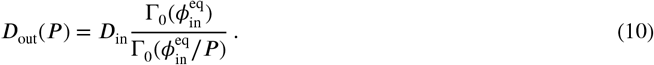

This expression indicates that for a given condensate with fixed 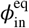 and ***D***_in_, there is a relationship between the diffusivity outside ***D***_out_ and the partition coefficient *P*. However, except for the limit *P* → 1, Eq. (10) does not impose further constraints on ***D***_out_(*P*) due to the unknown mobility function Γ_0_(*ϕ*_tot_). For large *P*, the missing knowledge of the mobility function renders ***D***_out_, ***D***_in_ and *P* as effectively independent parameters. This provides a theoretical justification for the phenomenological assumption given by (***Taylor et al. (2019))***, at least for large *P*.

In the following, we use our theory (Eq. (6)) to investigate the impact of the condensate environment on the FRAP dynamics. In particular, we consider how a passivated coverslip (no wetting of condensates) and nearby condensates affect the influx and thereby the recovery dynamics. Finally, we show how our theory can be used to determine the physical parameters corresponding to the condensate environment such as ***D***_out_ and *P* from a single FRAP experiment.

### Impact of non-wetting coverslip on FRAP dynamics

Here, we investigate the influence of the coverslip surface on the FRAP dynamics of non-wetting spherical droplets. Under typical *in vitro* conditions, condensates sediment due to gravity, leading to sessile droplets on a coverslip. In many experimental setups, coverslips are passivated, e.g. pegylated, in order to suppress wetting of condensates on the coverslip surface (***Alberti et al. (2018))***. These experimental conditions lead to almost spherical droplets since capillary effects are typically negligible for *μ*m-sized polymer-rich or protein-rich droplets (***Park et al. (2013); McCall et al. (2020))***.

We numerically solved Eq. (6) for a spherical condensate on top of a no-flux boundary that mimics the coverslip surface (Fig. 3a). We find that the recovery of the average volume fraction inside the condensate can slow down compared to the case without a coverslip (Fig. 3b). This slow-down vanishes if droplets have a distance to the coverslip surface larger than a few droplet diameters. Moreover, we find that the slow-down is more pronounced for larger partition coefficients *P*. This trend can be explained by the size of the region in the dilute phase from where most unbleached molecules are recruited: if *P* is small, most of the unbleached molecules come from the immediate surroundings of the condensate. Hence the influence of radial asymmetry is minimal and the recovery appears almost unchanged compared to the case without coverslip. However, for large *P*, the condensate recruits unbleached molecules from distances far away, limiting the recruitment to an effective half-space compared with the case without coverslip. Indeed, for very large *P* recovery rates slow down maximally by a factor of two.

**Figure 3.**
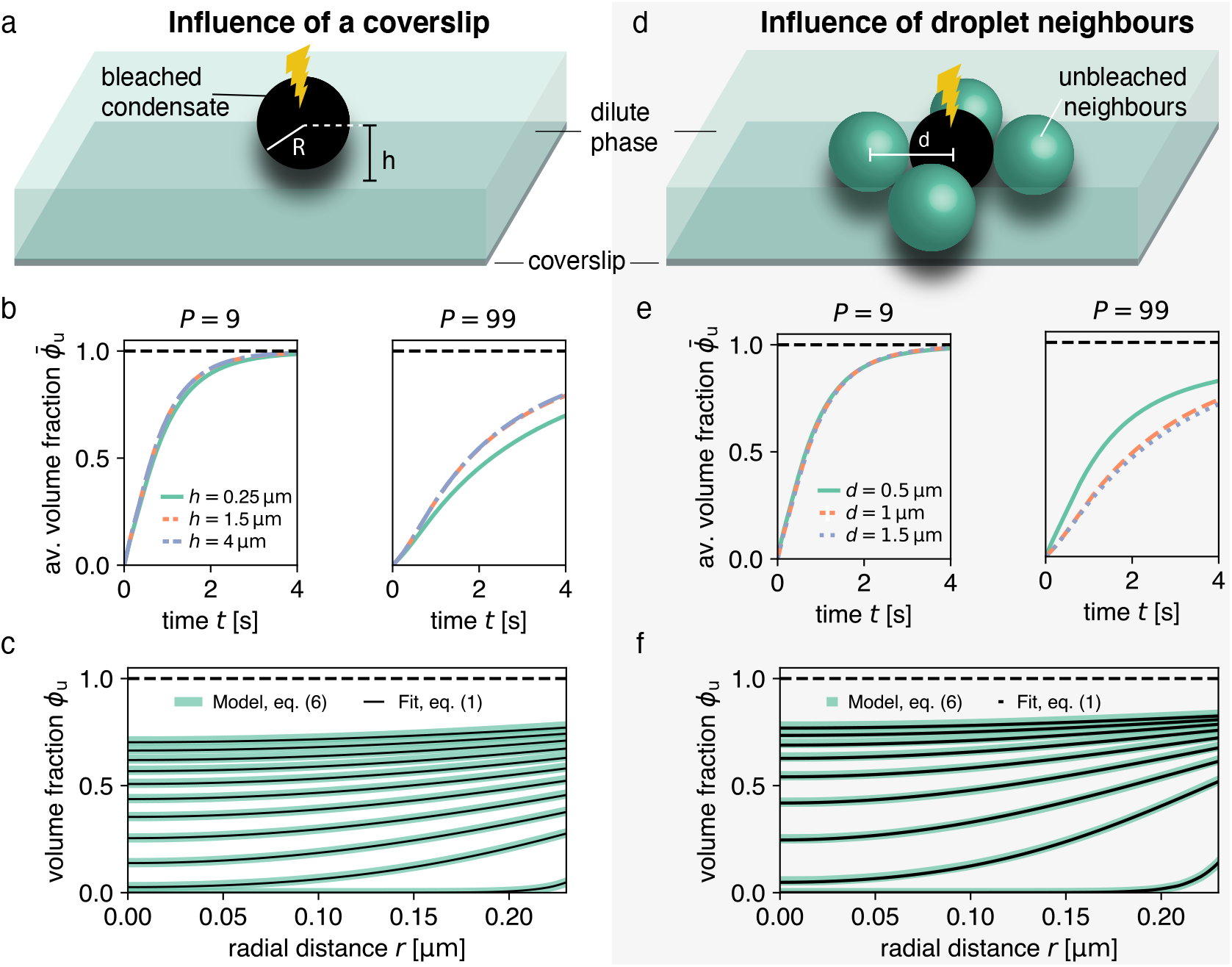
Impact of droplet environment on recovery dynamics. (**a**) Sketch of a typical experimental set-up with a droplet above a passivated coverslip, where the droplet center has a distance *h* > ***R*** to the coverslip. (**b**) Recovery of average unbleached volume fraction 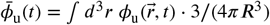 for different heights *h* above the coverslip at different partition coefficients ***P***. Results were obtained from finite element studies of Eq. (6) considering the geometries depicted in (a). For even larger *h*-values (e.g., no coverslip), results are approximately equal to the blue dashed line. (**c**) Using the method introduced in Fig. 1 on the worst case scenario *h* = 0.25 *μ*m in (a) results in an excellent fit and can reliably extract the input ***D***_in_. (**d**) Sketch of neighbouring droplets next to a bleached droplet. (**e**) Total recovery curves for finite element simulations of the geometry depicted in (d), for different distances between neighbouring droplet centers, at different partition coefficients ***P***. Note the strong dependence on the distance of neighbouring droplets. For even larger ***D***-values (e.g., no neighbouring drops), results are approximately equal to the blue dashed line. (**f**) Same as (c) but for worst case of (e), i.e. ***D*** = 0.5 *μ*m.

Interestingly, by extracting the boundary concentration in mid-plane, similar to the procedure in Fig. 1, and spatially fitting the solutions of Eq. (1) to the ensuing recovery profiles, we can reliably recover the input diffusion coefficient ***D***_in_ (Fig. 3c). The reason for this excellent agreement is that by considering the intensity at the condensate interface, our method is independent of the time scale set by diffusion in the dilute phase.

### Impact of neighbouring condensates on FRAP dynamics

In this section, we address the impact of neighbouring condensates on the recovery dynamics. We solved Eq. (6) for a system composed of a bleached condensate with four unbleached neighbouring condensates of the same size (Fig. 3d) and find that neighbouring condensates can significantly speed-up the recovery dynamics (Fig. 3e). This speed-up is only pronounced for rather close condensates with inter-droplet distance on the order of condensate size. Moreover, similar to the impact of the coverslip, the effects of the recovery dynamics are stronger for larger partition coefficients *P*. For small *P*, most unbleached molecules are recruited from the dilute phase leading to almost no effect on the recovery also when condensates are very close to each other (Fig. 3e, left). For large partition coefficients *P*, however, a certain fraction of molecules are recruited from the neighbouring condensates causing a significant speed-up of the recovery (Fig. 3e, right).

Again, despite this change in total recovery due to close-by neighbouring droplets, we can reliably measure ***D***_in_ via our spatial fitting method (Fig. 3f). In particular, by extracting the boundary intensity in mid-plane and spatially fitting the solutions of Eq. (1), we find very good agreement with our input ***D***_in_. This agreement shows that our method is robust for typical experimental systems that deviate from an ideal isolated condensate.

### How to determine partition coefficients and outside diffusivity via FRAP of condensates

We have shown that by using the time-dependent fluorescence at the interface of a spherical droplet we can accurately fit our dynamic Eq. (1) to our experimental data and thus determine the diffusion constant inside the droplet, ***D***_in_ (Fig. 1a). Our theory (see Eq. 6) suggests that the fluorescence at the droplet interface is affected by the physical parameters characterizing the droplet environment such as the diffusion coefficient ***D***_out_ and the partition coefficient ***P***. In particular, the flux through the droplet interface is enlarged for increasing ***D***_out_ or decreasing ***P*** (see Eq. (8c) after rescaling the concentration close to interface). Thus, for a condensate with concentration 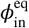 and diffusion coefficient ***D***_in_, the flux between both phases at the interface implies a relationship between ***D***_out_ and ***P***, as suggested by Eq. (10). However, due to the unknown mobility function, we need to obtain this relationship from experimental data.

Here, we determined the relationship between ***D***_out_ and ***P*** by fitting numerical solutions of Eq. (6) to the recovery dynamics inside the droplet (Fig. 4a-c and supp. movies 1,2,3). The diffusion coefficient inside, ***D***_in_, was independently determined for each experiment via our method introduced in Fig. 1a. This leaves ***P*** and ***D***_out_ as independent parameters, which is valid for large ***P*** (see discussion after Eq. (10)). We thus sampled ***P*** (with ***P*** ≫ 1) across three orders of magnitude and obtained the best-fitting ***D***_out_ for each ***P*** (Fig. 4d,e). Notably, all the combinations of ***D***_out_ and ***P*** represent relatively good fits. For very large ***P***, we find that the best ***D***_out_ scales linearly with ***P*** (dashed lines in Fig. 4d,e); for a discussion on the origin of this scaling, please refer to Appendix 2.

**Figure 4.**
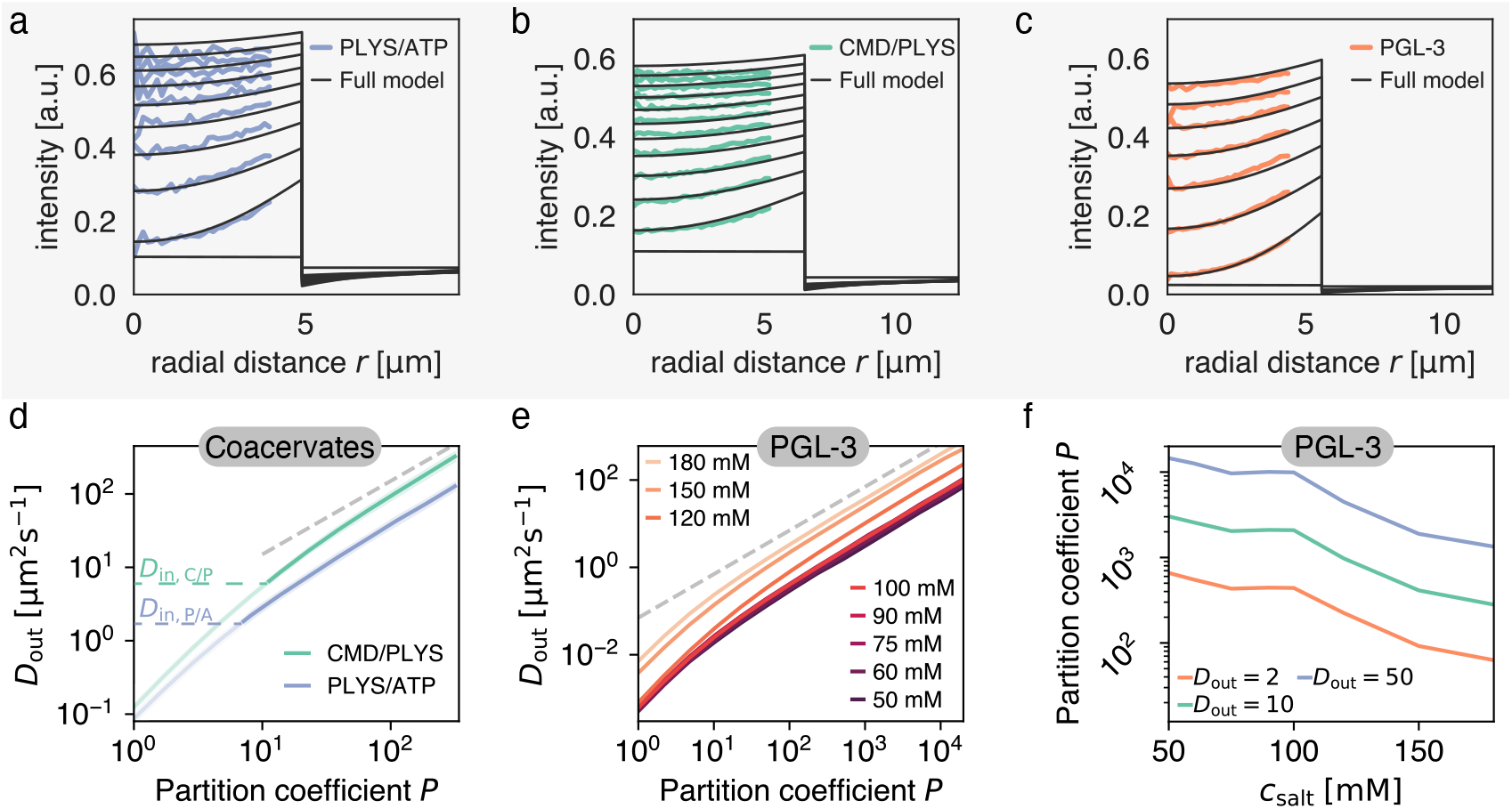
Varying partition coefficient P and diffusivity outside D_out_ simultaneously can lead to similar recovery kinetics. (**a**) Spatial recovery of a single PLYS/ATP droplet (blue) with a fit to the full model (black). Note, data close to the droplet boundary cannot be fit, due to optical artefacts giving rise to an artificially broad interface (see methods and Fig. 1a). (**b**) Same as (a) for a CMD/PLYS coacervate droplet. (**c**) Same as (a) for a PGL-3 droplet. (**d**) Given the partition coefficient ***P***, ***D***_out_ is found by fitting the coacervate data to the model. Note the convergence to a power law, ***D***_out_ ∝ ***P***^n^ with *n* =1 for large partition coefficients (grey dashed line; for a discussion see Appendix 2). Average ***D*** in values as obtained in Fig. 1f are indicated for both systems. Values with ***D***_out_ < ***D***_in_ are light shaded. (**e**) Same as (b) but for PGL-3 with different salt concentrations. Note the order from top to bottom from highest to lowest salt concentration. (**f**) Based on (e), partition coefficients ***P***(*c*_salt_) can be estimated for a given ***D***_out_. ***P***(*c*_salt_) is shown for different ***D***_out_. Units of ***D***_out_ in *μ*m^2^ s^-1^.

Strikingly, each experimental condition leads to a unique relationship between ***D***_out_ and ***P***. This relationship allows one to obtain the partition coefficient ***P*** by measuring ***D***_out_, e.g. by using FCS. We can also use this relationship to obtain insight into how the addition of salt affects the partition coefficient of PGL-3. The outside diffusivity ***D***_out_ can be estimated based on the value for diffusion of eGFP in water, ***D***_out,GFP_ = 87 *μ*m^2^ s^-1^ (***Arrio-Dupont et al. (2000))***. eGFP consists of 239 amino acids, while PGL-3 consists of 693 amino acids. If we assume the tagged protein to be spherical and use the Stokes-Sutherland-Einstein relationship ***D*** ∝ 1/*a*, we obtain ***D***_out_ ≈ 44 *μ*m^2^ s^-1^. Specifically, for salt concentrations in the range from 50 mM to 180 mM, the partition coefficient ***P*** of PGL-3 droplets decreases more than 10-fold (Fig. 4f). This trend is probably a result of enhanced screening of charged groups for increasing salt concentration and is in line with unpublished results using a technique relying on the measurement of droplet volumes (A.W. Fritsch and J.M. Iglesias-Artola, personal communication). To obtain more precise estimates of ***P***, Figs. 4d,e can be complemented by additional measurements of ***D***_out_. Note, the absolute values of ***P*** are overestimated up to twofold, since the fitting method employed here assumes spherical symmetry, thus effectively ignoring the effect of the coverslip (see caption of Fig. 3 for a discussion).

We next asked, whether there is a way to bypass this extra measurement and obtain specific values for ***P*** and ***D***_out_ at the same time, without measuring fluorescence intensities outside the droplet. Though each combination (***D***_out_, ***P***) along the lines specified in Figs. 4d,e leads to a reasonable fit, we will now show that there is a distinct combination that exhibits a global minimum of the cost function for each ***D***_out_(***P***). Providing experimental evidence of this global minimum can be hampered by environmental effects such as neighbouring droplets or the coverslip surface (Fig. 3a,d). In particular, in our experimental studies, interdroplet distances are sometimes on the order of the droplet size and diffusive exchange is affected by the coverslip. Thus, we decided to use our model to create *in silico* data and provide evidence for the existence of a distinct combination of ***D***_out_ and ***P*** for a fixed ***D_in_***. Fixing ***D**_in_* mimics the approach of initially determining ***D***_in_, as outlined in Fig. 1. To determine the relationship ***D***_out_(***P***), we proceed as described for Fig.4. Fig.5(a) depicts the ***D***_out_(***P***) relationships corresponding to four parameter combinations in a range relevant for protein condensates and coacervate droplets. In particular, we choose two outside diffusivities ***D***_out_ of 0.1 *μ*m^2^ s^−1^ and 1 *μ*m^2^ s^−1^ and two partition coefficients, ***P*** = 5 and ***P*** = 150. We find indeed that each cost function exhibits a unique minimum for each of the considered parameter combinations (Fig. 5(b,c)). These findings indicate that all three parameters, ***D***_in_, ***D***_out_ and ***P***, can in principle be determined by a single FRAP experiment of the droplet inside. Thus, in principle, there is no need to measure kinetic properties of the dilute phase to fully characterise the system in terms of its parameters. This possibility represents a new approach to determine the partition coefficient ***P***, which is particularly important in light of recent data showing that measurements based on fluorescence intensity can lead to drastic underestimation of ***P*** (***McCall et al. (2020)***).

**Figure 5.**
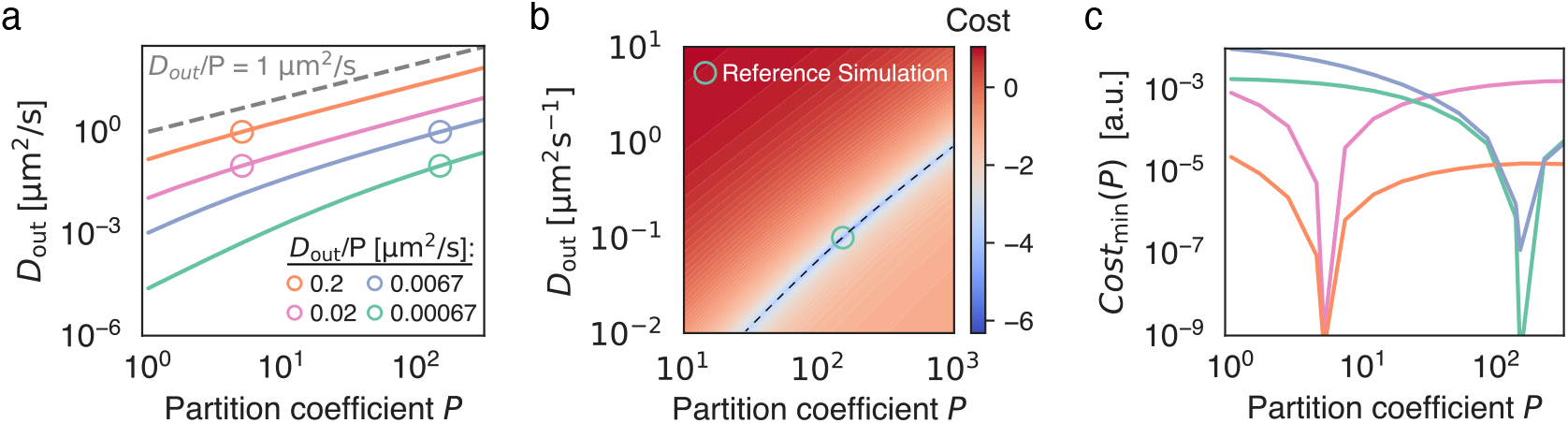
Given the recovery dynamics inside a condensate, the key parameters *D*_in_, *D*_out_ and *P* can be determined uniquely without measuring the outside dynamics. As reference systems, indicated by open circles, we consider *in silico* data, obtained by solving Eq. (6) with known parameters ***P*** and ***D***_out_. To mimic the approach of initially determining ***D***_in_(see Fig. 1) we keep ***D_in_*** = 0.01 *μ*m^2^ s^−1^ for all *in silico* datasets. (**a**) Given the partition coefficient ***P***, ***D***_out_ is found by fitting synthetically generated data to the model. (**b**) Cost function (colorbar, log-scale) as a function of ***D***_out_ and ***P***. We note that the minimum at the reference simulations coincides with the parameters used to generate the synthetic data. The valley in parameter space (dashed line) corresponds to the green line in (a). (**c**) Minimum of cost function for each ***P***, corresponding to curves shown in (a). This minimum corresponds to the valley indicated by the dashed line in (b). Note the minimum at the input parameter set, which indicates uniqueness of the outside dynamics for given values of ***D***_out_ and ***P***

## Discussion

The dynamic redistribution of fluorescent molecules has been used to characterize liquid phase separation in biology via a variety of techniques, including SPT, FRAP and FCS (***Elbaum-Garfinkle et al. (2015)***; ***Taylor et al. (2019)**; **Moon et al. (2019)***). Here, we have derived a theory from first principles that describes the diffusive motion of labeled molecules based on the physics of phase separation. It can be applied to many state-of-the art fluorescent methods such as FCS, SPT and FRAP and will thus be crucial help extend traditional techniques to the realms of phase separation ((***Ries and Schwille (2012)***)). Importantly, this theory enables us to avoid commonly applied approximations such as the frequently used single-exponential recovery (***Brangwynne et al.(2009)**; **Frottin et al. (2019***); ***Kaur et al. (2019***); ***Fisher and Elbaum-Garfinkle (2020***); ***Kistler et al. (2018***)).

Our theory describes the dynamics of labeled molecules through interfaces of condensates. It can be applied to spherical condensates but also non-spherical condensates and arbitrary bleach geometries, see Fig.3. We were able to quantify the impact of neighbouring droplets and the coverslip on the recovery dynamics. We found that neighbouring droplets caused an appreciable speed-up in overall recovery, while emulating a coverslip caused a weak slow-down. In order to experimentally verify our theory, we have used three *in vitro* droplet systems, two composed of charged synthetic polymers and one with a purified protein component. There is remarkable quantitative agreement between our theory and the diffusion dynamics observed inside such droplets. This agreement shows that proteins and charged, synthetic polymers form droplets that follow simple diffusive dynamics inside. Crucially, we use the full spatio-temporal data for fitting and can thus distinguish the timescale set by intra-droplet diffusion from the timescales at play in the dilute phase. We extract the intensity directly at the inside of the droplet interface and fit a spatially resolved diffusion equation to the ensuing recovery. We use the boundary intensity as a dynamic boundary condition and the experimentally measured profile as initial condition. Throughout the time course, we find excellent agreement with the data and have thus found a method with minimal approximations that can precisely measure the inside diffusion coefficient ***D***_in_.

Building on the analysis of the droplet phase, we show that there is a relationship between partition coefficient ***P*** and the outside diffusion coefficient ***D***_out_. Data obtained from FRAP experiments define a line in (***D***_out_, ***P***) space, along which any parameter set can reliably account for the boundary dynamics. This unique relationship allows measuring ***P***, when ***D***_out_ is known, and vice versa, which opens an interesting avenue for measuring partition coefficients purely based on dynamics. This is particularly important in light of recent data obtained by quantitative phase microscopy (QPM), which show that measuring partition coefficients based on fluorescence intensity can lead to strong underestimation of ***P*** (***McCall et al. (2020***)). In the future, it would thus be interesting to use FCS to determine ***D***_out_ and compare the partition ***P*** inferred by our method with measurements by QPM.

Our approach can be readily extended to multi-component systems with an arbitrary number of components, which is particularly useful *in vivo*. This would be hardly possible for techniques that do not use labeled components, such as QPM or other scattering methods. Of particular interest are multi-component systems with chemical reactions away from equilibrium. Our approach can then be used to determine the diffusion coefficients and concentration levels of reactants, and thereby provide insights into reaction kinetics. Interestingly, introducing the bleached molecules via a ternary mixture also enabled us to derive the Langevin equation governing single-molecule motion in phase-separated media, thus providing a link to SPT (***Bo et al. (2021***), in preparation). Approaches for single labelled molecules are highly relevant since high labeling fractions were shown to alter the kinetics in dense protein phases (***Jawerth et al. (2018, 2020***)). Finally, our techniques can also be employed to characterize rheological properties of condensates such as the recently reported glass-like dynamics of protein droplets (***Jawerth et al. (2020)***).

## Acknowledgements

We are grateful to P. McCall, A. Fritsch, J.M. Iglesias-Artola, G. Bartolucci, T. Wiegand, M. Karnat, E. Filippidi, T. Harmon, F. Jülicher and members of the Weber and Hyman groups for stimulating discussions, and P. McCall and J. Pfanzelter for very valuable and insightful comments on the manuscript. L. Hubatsch and C. Weber acknowledge the SPP 2191 “Molecular Mechanisms of Functional Phase Separation” of the German Science Foundation for financial support.

## Supplementary

**Movie S1: FRAP dynamics in a PGL-3 droplet**

Left: Representative *in vitro* droplet after full bleach. Time course starts after a small time lag due to a fast uniform recovery (see methods).

Middle: Diffusion eq. (1) fit to azimuthally averaged droplet intensity with a global fit parameter ***D***_in_.

Right: Full model eq. (6) fit to azimuthally averaged droplet intensity.

**Movie S2: FRAP dynamics in a CMD/PLYS coacervate**

For description see Movie S1.

**Movie S3: FRAP dynamics in a PLYS/ATP coacervate**

For description see Movie S1.

## Methods

### Coacervate assay

#### General reagents

Carboxymethyl-dextran sodium salt (CM-Dex, (C6H10O5)n.(COOH), 10-20kDa, monomer MW =191.3 gmol^−1^), Poly-L-lysine hydrobromide (PLys, (C6H12N2O)n, 4-15 kDa, monomer MW =208.1 g mol^−1^) and adenosine 5’-triphosphate disodium salt hydrate (ATP, C10H14N5Na2O13P3, MW =551.1 g mol^−1^) were purchased from Sigma Aldrich. FITC-PLys ((C6H12N2O)n.(C21H11NO5S), 25 000 g mol^−1^) was purchased from Nanocs, NewYork, USA. Milli Q water was used to prepare aqueous stocks of CM-Dex (1000 mM, pH 8), PLys (200 mM, pH 8) and ATP (100 mM, pH 8). All solutions were stored in the freezer at −20 °C until use and the pH of all stocks was adjusted using a stock solution of 1 M NaOH.

#### Coacervate preparation

Stock solutions of CM-Dex, PLys and ATP were first diluted to 25 mM and the PLys solution doped with 1% v/v PLys-FITC. Diluted solutions of CM-Dex/PLys or PLys/ATP were then mixed together at a 4:1 volume ratio (16 *μ*l: 4 *μ*l), resulting in the formation of turbid coacervate solutions. Solutions were left to equilibrate for at least 5 min before imaging, up to a maximum of 15 min when larger droplets were desired.

### PGL-3 droplets

PGL-3 was purified and stored as previously described (***Saha et al. (2016)***). To obtain droplets, 300 mM KCl stock protein solution was diluted to the desired concentration, achieving final salt concentrations of 50-180 mM. A small imaging volume was created by using polystyrene beads, resulting in complete droplet sedimentation after less than five minutes. Droplets were imaged immediately to avoid changes in material properties due to ageing (***Jawerth et al. (2020***)).

### Microscopy and FRAP

#### Confocal imaging

Droplets were imaged at midplane by visually defining the focal position with the largest droplet area of the droplet of interest. Images were acquired on an Andor spinning disk confocal microscope equipped with an Andor IX 81 inverted stand, a FRAPPA unit, an Andor iXON 897 EMCCD camera, and a 488nm laser, using a 60x/1.2 U Plan SApo OLYMPUS water objective. Imaging conditions were optimized for minimal bleaching at the required frame rate. Frame rates were optimised for each system: PGL-3, 0.1 s Δ*t* 5 s, CMD/PLYS, Δ*t* = 0.03 s, PLYS/ATP, Δ*t* = 0.07 s.

#### FRAP

Droplets were bleached in their entirety by using the minimal FRAP ROI that encompasses the entire droplet. FRAP was performed in three focal planes, equally spaced across the droplet in z-direction, to reduce non-uniform bleaching of the droplet. FRAP rates and dwell times were chosen such that left-over fluorescence intensity above background was smaller than 1% for PGL-3 and smaller than 15% for coacervate droplets to maximize bleaching within the droplet while keeping bleaching impact on the droplet environment minimal.

### Data analysis

#### Azimuthal averaging and normalization

Time-lapse images were cropped with the droplet of interest in the center. An azimuthal average was performed around the center of the droplet to obtain a 1D profile along the radial coordinate with minimal loss of data, using the *radialavg* function provided by David J. Fischer on Matlab File Exchange. Camera background was subtracted uniformly from the resulting 1D profiles. The radial intensity profile at the prebleach stage was used for normalization and to correct for optical artefacts that lead to increased fluorescence at the droplet center compared to the droplet-bulk interface. Data close to the droplet interface cannot be used for fitting, since the droplet has an artificially broad boundary due to the point-spread function and likely due to curvature effects. Therefore, on average, the intensity of the ten pixels closest to the boundary were not used for analysis. The droplet boundary was defined as the inflection point of the azimuthally averaged profile in the pre-bleach frame.

Immediately after bleaching a uniform recovery across the entire droplet can be seen, which cannot be spatially resolved even at frame rates < 30 ms. This recovery is fast compared to the recovery by diffusion from the outside for all systems under investigation. We thus chose to not account for this uniform recovery in our model and instead start the fitting after a time lag that depends on the system and droplet size. This offset typically consists of less than 5% of the total pre-bleach intensity. Additionally, bleaching is not complete, resulting in an additional offset above the camera background even immediately after bleaching (see grey lines in Fig.1).

Photo-bleaching due to continuous imaging was minimal in all droplet types. We thus chose to not account for imaging-induced photo-bleaching, in order to not introduce additional noise due to necessarily occurring fluctuations within the bleach correction.

#### Extracting experimental boundary conditions

*c_u_*(*r* = ***R***_, *t*) was extracted from the intensity profiles as the value at the outermost pixel. In order to speed up fitting and avoid jumps in *c_u_*(*r* = ***R***_, *t*), the extracted intensity values were sorted to eliminate small fluctuations.

Fitting of ***D***_in_ by using experimentally measured boundary conditions (Fig.1)

The resulting spatio-temporal profiles were fit as described in the main text, using ***D**_in_* as a single global fit parameter and using *c_u_*(*r* = *R*_, *t*) as described above as the system’s time-dependent boundary condition. Fits were performed in MATLAB (Mathworks), using pdepe to solve the PDE and fminsearch for minimizing the squared distance between data and model. Code is available at https://gitlab.pks.mpg.de/mesoscopic-physics-of-life/DropletFRAP.

### Numerical solution of Eq. (6)

Eq.6 was solved using either pdepde (MATLAB (Mathworks), Figs.4, 5, for spherically symmetric systems) or by using the finite element method via the FENICS environment (***Logg et al. (2012***)) for arbitrary 3D geometries (Fig. 3). All fits in Figs. 4,5 were performed using fminsearch based on a squared-difference metric. Code is available at https://gitlab.pks.mpg.de/mesoscopic-physics-of-life/frap_theory.

## Appendix 1

### Limit of narrow interfaces

Here we derive the effective droplet model for our dynamic equation

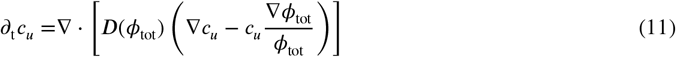

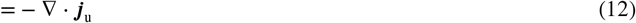

by considering the limit of narrow interfaces. In the equation above, ***J***_u_ denotes the flux of unbleached molecules. Conservation of molecules at the interface implies

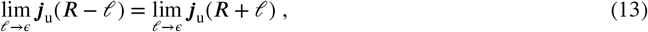

where *ŕ* is the characteristic size of the interface. Moreover, *ϵ* > 0 is a small but non-zero parameter by which we define the position directly left and right of the interface. In this limit, ∇*φ*_tot_|_***R***±*ϵ*_ = 0 directly left and right of the interface. Thus, we obtain Eq. (8c) for narrow interfaces. However, the contribution ***D***(*φ*_tot_)*c_u_*∇*φ*_tot_/*φ*_tot_ in Eq. (11) that determines the dynamics through the interface implies a Dirichlet boundary conditions for the concentration of unbleached molecules at the interface *r* = ***R***. Since ∇*φ*_tot_/*φ*_tot_ ∝ ŕ, we demand

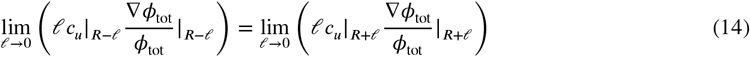

in the limit of decreasing interface width *ŕ*. This condition ensures that the dynamics at the interface remains unchanged for decreasing *ŕ*. Parameterizing the interface for example by 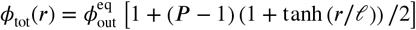, we find that Eq. (14) leads to Eq. (8d).

## Appendix 2

### Solution of effective droplet model for FRAP

In this appendix we derive the solution for the two diffusion equations coupled at the interface for a spherically symmetric case; see Eqs. (8). We consider an initial condition with a constant concentration *c*(***r***, 0) = *c*_0_ outside a droplet with radius ***R*** and bleaching gives rise to *c*(***r***, 0) = 0 inside the droplet. Performing a Laplace transformation, 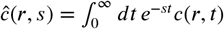, of Eqs. (8), we find

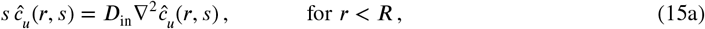

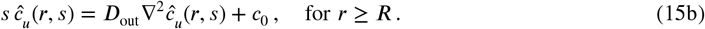

where *s* is the rate parameter of the Laplace transform. The corresponding solutions read

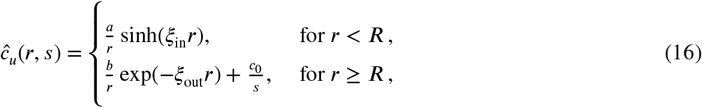

where 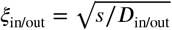. We have selected the solutions with no radial flux at the origin and a finite concentration at infinity. The remaining unknown constants *a* and *b* are determined by the conditions at the interface stated in Eqs. (8). Inside the droplet *r* < ***R***, the solution reads

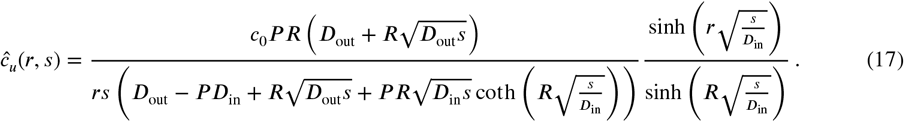

Even though it is non-trivial to perform the inverse Laplace transformation of this expression we can obtain some analytical understanding of the behaviour at short and long timescales by considering the asymptotics of large and small *s*. Expanding Eq. (17) for small *s*, we obtain:

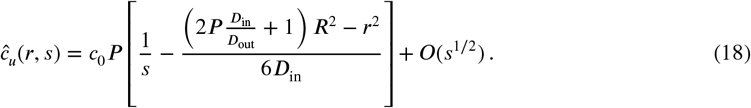

This relationship shows that for small *s*, the rescaled concentration *ĉ*_u_(***r**, s*)/***P*** is affected by ***D***_out_ only via the ***PD***_in_/***D***_out_ combination. Conversely, for large *s*, the leading contribution features the term

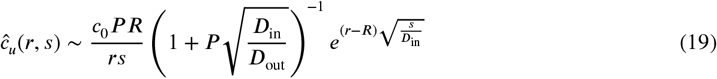

showing that for short times the system is influenced by ***D***_out_ via the ratio 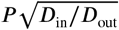. Note that for these short times the solution is well approximated by that of a one-dimensional system. These long and short time behaviors are confirmed by the numerical solutions plotted in Appendix 2 Fig. 1, which show that the evolution of the rescaled concentrations match at short times, if 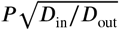 is kept constant but start to deviate for longer times. Conversely, at longer times the dynamics are invariant if ***PD***_in_/***D***_out_ is kept constant except for a shift due to a short timescale transient.

**Appendix 2 Figure 1.**
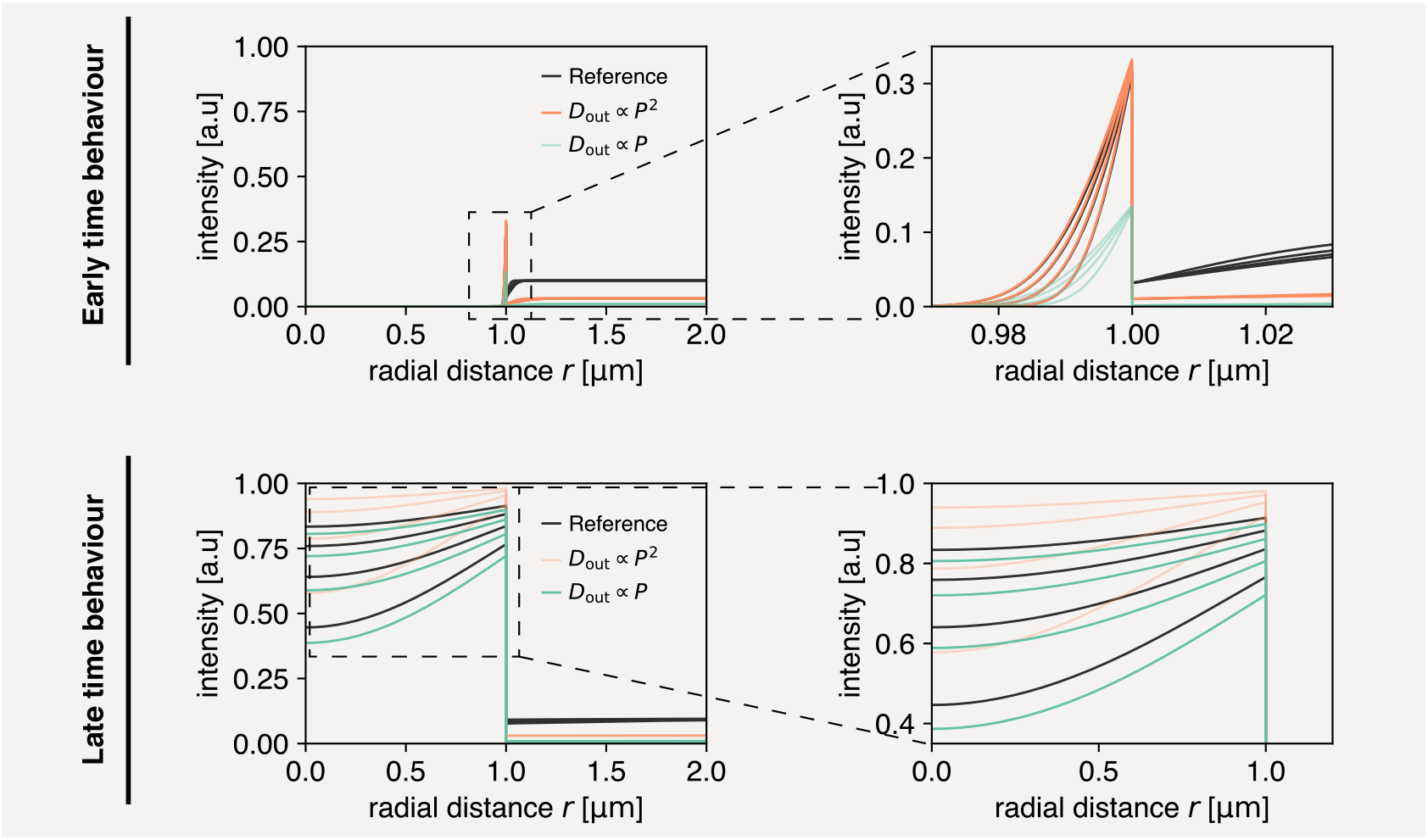
Solution to dynamics inside droplets is unique, but early and late time behaviour can be approximated by different scaling regimes. **(Top)** Early time behaviour of numerical solutions. If a parameter set is chosen (‘Reference’), at early times this can be approximately recovered by scaling ***D***_out_ ∝ ***P***^2^ (orange). A linear scaling does not recover the same inside dynamics at early times (green). **(Bottom)** Late time behaviour of numerical solutions. At late times the reference dynamics can be approximately recovered by scaling ***D***_out_ ∝ ***P*** (green). Meanwhile, ***D***_out_ ∝ ***P***^2^ does not recover the reference inside dynamics (orange).

## Notes

https://gitlab.pks.mpg.de/mesoscopic-physics-of-life/DropletFRAP

https://gitlab.pks.mpg.de/mesoscopic-physics-of-life/frap_theory

### Competing Interest Statement

The authors have declared no competing interest.

